# StrongestPath: a Cytoscape application for protein-protein interaction analysis

**DOI:** 10.1101/2020.11.12.380600

**Authors:** Zaynab Mousavian, Mehran Khodabandeh, Ali Sharifi-Zarchi, Alireza Nadafian, Alireza Mahmoudi

**Author notes:** **Corresponding Author**, Zaynab Mousavian, Ph.D, Department of Computer Science, School of Mathematics, Statistics, and Computer Science, University of Tehran, Tehran, Iran.

## Abstract

**Background:** StrongestPath is a Cytoscape 3 application that enables to look for one or more cascades of interactions connecting two single or groups of proteins in a collection of protein-protein interaction (PPI) network or signaling network databases. When there are different levels of confidence over the interactions, it is able to process them and identify the cascade of interactions having the highest total confidence score. Given a set of proteins, StrongestPath can extract and show the network of interactions among them from the given databases, and expand the network by adding new proteins having the most interactions with highest total confidence to the current proteins. The application can also identify any activation or inhibition regulatory paths between two distinct sets of transcription factors and target genes. This application can be either used with a set of built-in human and mouse PPI or signaling databases, or any user-provided database for some organism.

**Results:** Our results on 12 signaling pathways from the NetPath database demonstrate that the application can be used for indicating proteins which may play significant roles in the middle of the pathway by finding the strongest path(s) in the PPI or signaling network.

**Conclusion:** Easy access to multiple public large databases, generating output in a short time, addressing some key challenges in one platform and providing a user-friendly graphical interface make the StrongestPath easy to use.

## Background

The teamwork of proteins, in terms of temporary or permanent interactions, is critical for the majority of the biological processes. So far there have been numerous protein-protein interaction (PPI) or signaling pathways databases developed based on the experimental approaches or computational predictions. Some of the databases assign different confidence levels to the interactions, e.g. higher confidences for the experimentally validated interactions, and lower values for the computationally predicted ones. So, the identification of the cascades of interactions from the receptors to the transcriptional regulatory factors is a major challenge in systems biology. Cytoscape [1, 2] is a highly flexible platform for integrating new pieces of software called Apps, and there are a number applications for Cytoscape to compute paths in the biological networks like PesCa [3], PathExplorer (http://apps.cytoscape.org/apps/pathexplorer) and PathLinker [4, 5].

We developed StrongestPath, a Cytoscape 3.0 application to address a number of key challenges during analysis of PPI or signaling networks. The first challenge is identifying a cascade of interactions, as a regulatory or signaling pathway, in a large PPI or signaling network. In many biological studies perturbation of a protein *A* is observed to influence a protein *B,* but the cascade of interactions between *A* and *B* is unidentified. The number of possible pathways of any length between *A* and *B* can be extremely high, and finding a short pathway with highly confident interactions is not straightforward. The problem will be more complicated if there are two distinct groups of proteins, and the target is finding short and confident pathway having one end in each group. The network interactions can be indirect, directed or mixed, and in either case StrongestPath indicates short and highly confident pathways. It can return optimal pathways or a collection of sub-optimal pathways having total confidence above a user-defined threshold with possible overlaps.

The second challenge addressed by StrongestPath is growing the sub-network of some proteins, either by extracting their pairwise interactions from a list of PPI or signaling databases, or adding further proteins that are more likely to create protein complexes or dense interactions with existing proteins. For this purpose, StrongestPath looks the whole PPI or signaling network and identifies proteins with maximum total confidence of interactions to the given set of proteins. This feature can be used for identification of unknown elements of a protein complex, biological process or core regulatory circuitry. Moreover, the third challenge addressed in the application is answering the question of whether there is any activation or inhibition regulatory path between two distinct groups of proteins. For example, when a list of genes is identified in a study of a phenomenon, researchers seek to answer the question of whether there is a regulatory pathway between the transcription factors associated with the phenomenon and the identified genes. In the Regulatory Path panel of the StrongestPath application, the application searches the TRRUST database [6], a manually curated transcriptionally regulatory network, to answer such a question.

StrongestPath comes with two types of built-in databases: (I) some PPI and signaling networks of human and mouse, containing interactions recorded in public databases, (II) protein nomenclature database, containing 11 different symbols and accession IDs of genes and proteins in different databases. Although more species can be added to the application in the future without any new installation, currently users can provide their own networks and nomenclature datasets, which allows StrongestPath to be used for any organism, PPI, and also gene regulatory and signal transduction networks.

Our results on 12 signaling pathways from the NetPath database indicates that the idea of identifying strongest path can be helpful for pathway reconstruction, and since the stored interactions in different databases may vary, simultaneous search of multiple databases can be a good solution. Among the available Cytoscape apps, the PathLinker is more similar to ours in terms of functionality and it was shown in [4, 5] that the PathLinker performs better than several state-of-the-art algorithms in pathway reconstruction. Therefore, we only compare our application with the Pathlinker in the results and discussion part of the manuscript. Easy access to multiple public large databases, generating output in a short time, addressing some key challenges in one platform and providing a user-friendly graphical interface make the StrongestPath easy to use.

## Implementation

The StrongestPath is designed with four main panels including *select databases, strongest path, expand* and *regulatory path.* In the following, we describe each section separately:

### Select Databases

We developed StrongestPath in Java, along with R scripts to preprocess the required databases. We used the NCBI [7] and the UniProt [8] databases to build the built-in protein-coding genes nomenclature databases, which allows to use any of 11 different gene or protein accession numbers including Entrez Gene ID, Official gene symbol, Aliases, Uniprot Gene ID, Ensembl (gene, transcript and peptide), Ref-Seq (peptide and mRNA), Reactome ID and STRING ID. We also supplied the application with some PPI, signaling and regulatory networks from public databases including STRING [9], HitPredict [10], HIPPIE [11] (only for human), KEGG [12], Reactome [13] and TRRUST [6]. Currently, both human and mouse species are supported in the application. However, the application can be easily expanded for supporting more species and also networks from more public databases in the future without having to install another version of the application and only with internet connection.

Whenever the user start the application, if the internet connection would be available, the list of supporting species by the application will be updated and the user can easily and quickly access to the all available databases for the selected species by only clicking on the *Download/Update Databases* button. We retrieved network data from public databases and made new network data with very smaller size according to the built-in nomenclature database. Currently, the downloaded data can be stored in hard disk drive with less than 1GB free space and the application can be used later without any dependence on internet connection.

Furthermore, users can use the application with their own data including the annotation file and the network file. In the annotation file, the user can provide different types of identifiers for the network nodes by a text file with multiple columns separated by Tab. In this file, each row refers to a specific node of the network and each column represents a list of a specific identifier type, separated by “,”. Only the first column of the annotation file, which is used to display nodes, is required and additional columns are optional in this file that allow users to access nodes in the application with multiple accession identifiers. The network file is a three-column text file, separated by Tab, in which each interaction is reported in one row and the columns refer to the source node, the target node and the confidence score (i.e. a probability value between 0 and 1) respectively. All of the nodes identifiers given in the annotation file can be used in the network file.

The StrongestPath application is developed to work with the current version of the Cytoscape. As mentioned earlier, it implements different scenarios in three distinct panels of *Strongest Path, Expand* and *Regulatory Path.* In each run of the application, firstly the built-in databases of the selected species or the user provided data should be loaded in the application by pressing the *Loading Databases* button in the *Select Databases* panel and then the application can be followed in other panels (See Fig.1).

**Fig. 1.**
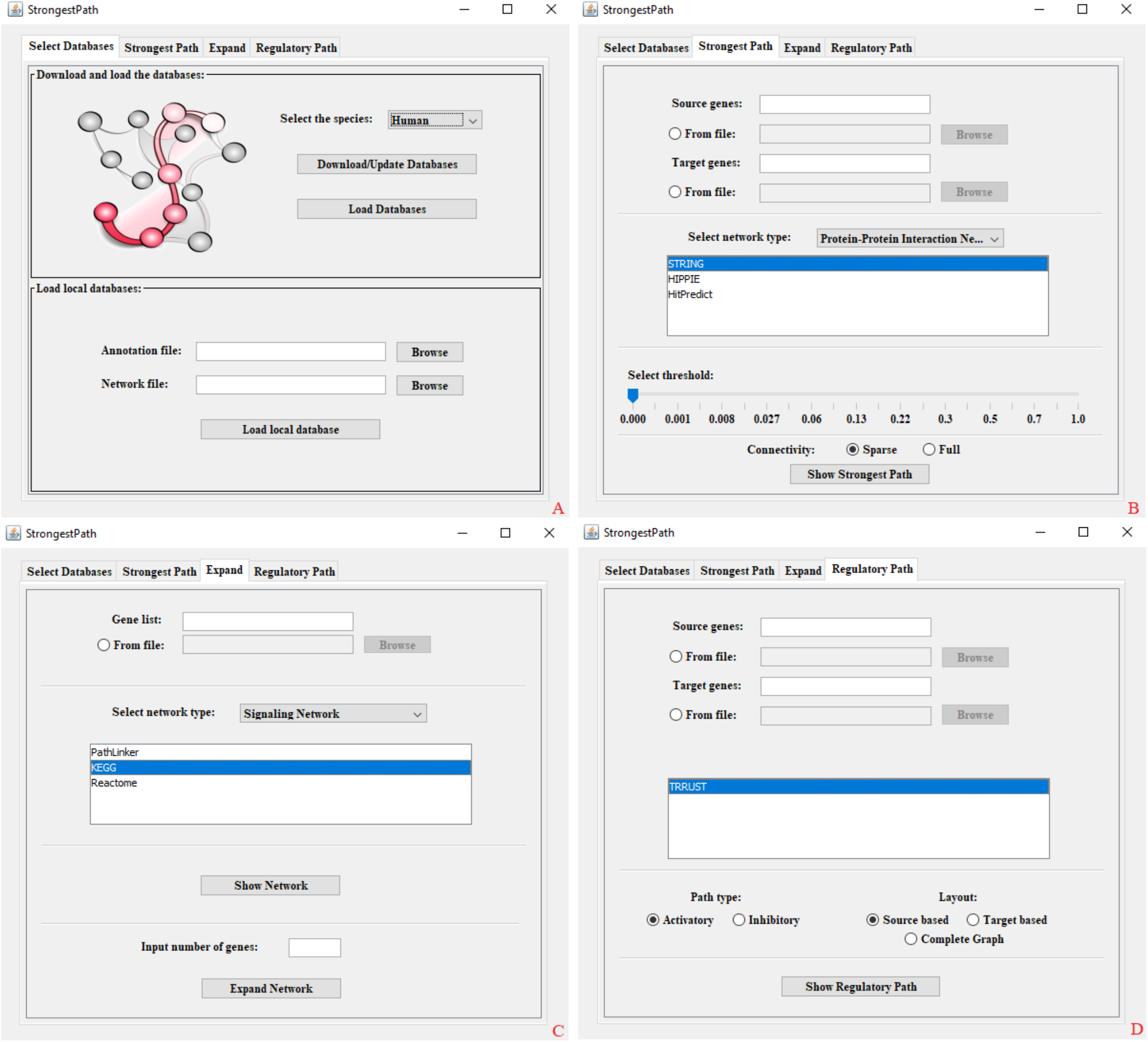
A view of StrongestPath with four panels: A) Select Databases, B) Strongest Path, C) Expand and D) Regulatory Path

### Strongest Path

We used interaction confidence scores for assigning weights between 0 and 1 to each edge of the PPI or signaling network. Given two sets of proteins, our goal is to identify the strongest path connecting at least one protein of the first set to one protein of the second. Thinking confidence scores as the probability of interaction, we define the strongest path as most probable chain of interactions, i.e. having maximum product of the edge probabilities. In different networks, the strongest path between specific nodes of network has a different meaning. While the strongest path can represent the most likely chain of interactions between two groups of proteins, it also represents a linear signaling pathway while the given graph is a signaling network [14, 15].

It is easy to show that identifying the path between two nodes of a general graph with maximum edge weights product is NP-Complete: by assigning a constant weight 2 to every edge, the problem would be equivalent to the HAMILTONIAN PATH, which is also NP-Complete. However, we made a “dual” graph with the same set of edges as the original one but with modified weights, using it the exact solution can be found in polynomial time. It is only required that all original weights are positive numbers not greater than 1, which is the case while the original weights are interaction probabilities.

Here is the formal problem statement: We have a weighted directed or in-directed graph *G* with all weights as positive real numbers not greater than 1, that we call it “primal” graph. Two disjoint subsets of nodes *A* and *B* are also given from the nodes of *G.* We are looking for a path *y* from the set of all possible paths *S* from any node in *A* to any node in *B*, having maximum weights product:

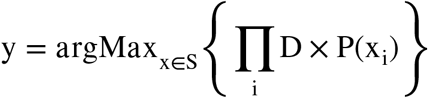

Where *x_j_* is the *i*-th edge of a path *x*, *P*(*x_i_*) is the interaction probability assigned to that edge, and *D* is a constant penalty factor on the lengths of the paths. In the dual graph, we change the weight of each interaction (edge) *e* by *w*(*e*) = – (log*D*+log(*P*(*e*)). Hence we can find the strongest path *y* as:

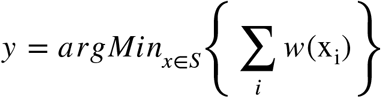

Having 0 < *P*(*e*) ≤ 1 for all edges *e* and *D*=0.95, the -(log*D* + log(*P*(*e*)) would be always positive non-infinite real values. Hence, we can apply a normal shortest-paths algorithm (i.e. Dijkstra’s algorithm) on the dual graph. Our method works not only for finding the strongest path between two single nodes, but also two groups of nodes.

We also generalized the algorithm to find sub-optimal strongest paths (those having slightly less probability product than the maximum). For a given positive real value *ϵ*, we define *ϵ*-strongest paths as follows:

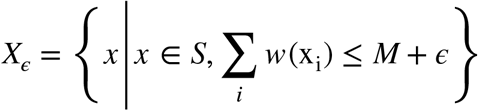

Where *M* is the length of the optimum shortest path in the dual graph:

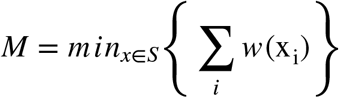

To find out *X_ϵ_*, again we use the dual graph and for every node *v*, we define *a*(*v*) as the weight of shortest path from any node of *A* to *v*, and similarly *b* (*v*) as the shortest weight to reach from *v* to any node in *B*. Pruning all nodes having *a*(*v*) + *b*(*v*) > *M* + *ϵ* can be shown to leave a graph T having exactly all nodes in *X_ϵ_*. The number of paths in *X_ϵ_* can exponentially grow with increased *ϵ*, however finding the proteins in the middle that can play a role in chain of interactions between *A* and *B* is suitable for most of applications. For better visualization of result, the Breath First Search (BFS) algorithm is employed to compute for each node the depth from *A*. The color and position of the nodes are then assigned accordingly.

Using the *Strongest Path* panel in the application, the user can find the strongest path connecting at least one source protein to one target protein, as described above. A comma-separated list of source genes and target genes can be given as input to the application by entering any accession gene identifier supporting in the application. As seen in Fig. 1, list of genes can also be given to the application via a text file containing one line per gene. By choosing a network type from one of the default types, a list of networks of that type, supporting in the application, is displayed for selection. The *Show Strongest Path* button searches for the strongest path between the source and target nodes in the selected networks, and the output of each selected network is shown in Cytoscape as a separate network. Before any search, at least one source gene, one target gene and one network must be selected by the user. As seen in Fig. 1, a ruler is provided at the bottom of the panel. The user can find the sub-optimal strongest paths between source and target nodes, as defined above, by increasing the threshold value defined on the ruler. When increasing the threshold parameter, the number of strongest paths increases exponentially and the output will be dense. By selecting the sparse option, although all proteins that appear on at least one strongest path are displayed, all strongest paths are not displayed, which saves running time and makes the output network sparse.

### Expand

In this panel, a list of input proteins is given to the application and the application returns a network containing input proteins and their connections in the selected background network at the first step. When giving an input positive integer *n*, the network is expanded by adding *n* proteins whose total weight of interactions with the proteins in the network has the highest value. A list of input genes can be entered into the application directly or by providing a text file. The input format in the whole application is similar to what was mentioned earlier. After choosing the network type, a list of loaded networks in the application, is given for selection as indicated in Fig. 1. The *Show Network* button searches for a network of interactions among input genes based on the interactions reported in the selected databases. If the user select more than one database, the network associated with each database is shown in Cytoscape separately. Each of these networks can be expanded by a number of close neighbours by clicking the *Expand Network* button.

### Regulatory Path

To answer whether there is any activation or inhibition path between source genes, encoding transcription factors, and target genes, the user can use the *Regulatory Path* panel (Fig. 1). We used the BFS algorithm to compute the shortest path between any pair of source and target genes. In this case, we defines the shortest path as a path connecting source and target genes with the minimum number of links. If the path is available, weights of +1 and −1 are assigned to activation and inhibition links of the path respectively, based on the information about mode of regulation in the experimentally verified databases like TRRUST [6]. In the simplest case, there are two situations for each regulatory path, when a change of level of source gene causes to the change of level of target gene. If the presence of source gene implies the presence of target gene and conversely the absence of source gene implies the absence of target gene, the path is called activation path. The opposite situation corresponds to the inhibition path. Accordingly, if the edge weights product is +1, the shortest path is defined as an activation path, otherwise the path is defined as an inhibition path.

In this panel, source and target genes can be inputted to the application as same as the other panels. Currently, only TRRUST database is available in the application for finding regulatory paths, but more databases would be added to the application in future. TRRUST is an experimentally validated database containing human TF-target links with mode of regulation information. It is worthy to note that the user can quickly access to any available database in each run of the application whenever they press the *download/update databases* button in the first panel. After pressing the *Show Regulatory Path* button, the application computes the regulatory paths with the selected mode of regulation between source and target genes, as defined above, based on the reported information in the selected database.

## Results

To demonstrate the application of StrongestPath, we used 12 signalling pathways provided in the Net-Path [16] database. The signalling receptors and transcription factors of each pathway were identified using the NetSlim [17] and the MSigDB [18] databases respectively. For each pathway, the receptors and TFs were given as source and target genes to the application respectively, and then we used the application to find the strongest path(s) between sources and targets in a background network. Since two types of networks including signalling networks and protein interaction networks can be selected in the application as a background network, we selected KEGG and STRING networks in separate application runs. The current version of KEGG network, derived from the aggregation of all KEGG signalling pathways, includes 6326 proteins and 61980 interactions, and since KEGG is a curated database, the probability score of all network interactions were considered equal to 1. The STRING network is a very large protein interaction network consisting of 18725 proteins and more than 5 million interactions, and all links were weighted by a confidence score between 0 and 1. Although the background networks, specially the STRING network, are very large, the application is able to find the strongest path(s) between multiple source and target genes in less than 10 seconds. In addition of strongest path(s), we also identified sub-optimal strongest path(s) between source and target genes in the STRING network by increasing the threshold parameter three times. Since all links of the KEGG have the same probability score, the number of links in the path determines the weight of the path and increasing the threshold parameter, in most cases, leads to the addition of a large number of genes to the detected sub-network. Therefore, we used the application to identify only the strongest path(s) between source and target genes in the KEGG network containing at least one gene in the middle of the path. To assess the performance of the application, for each pathway, we investigated how many of the genes found by the application in each run were already known as pathway genes in the NetPath database. The obtained results are given in Tables 1 and 2.

**Table 1.**
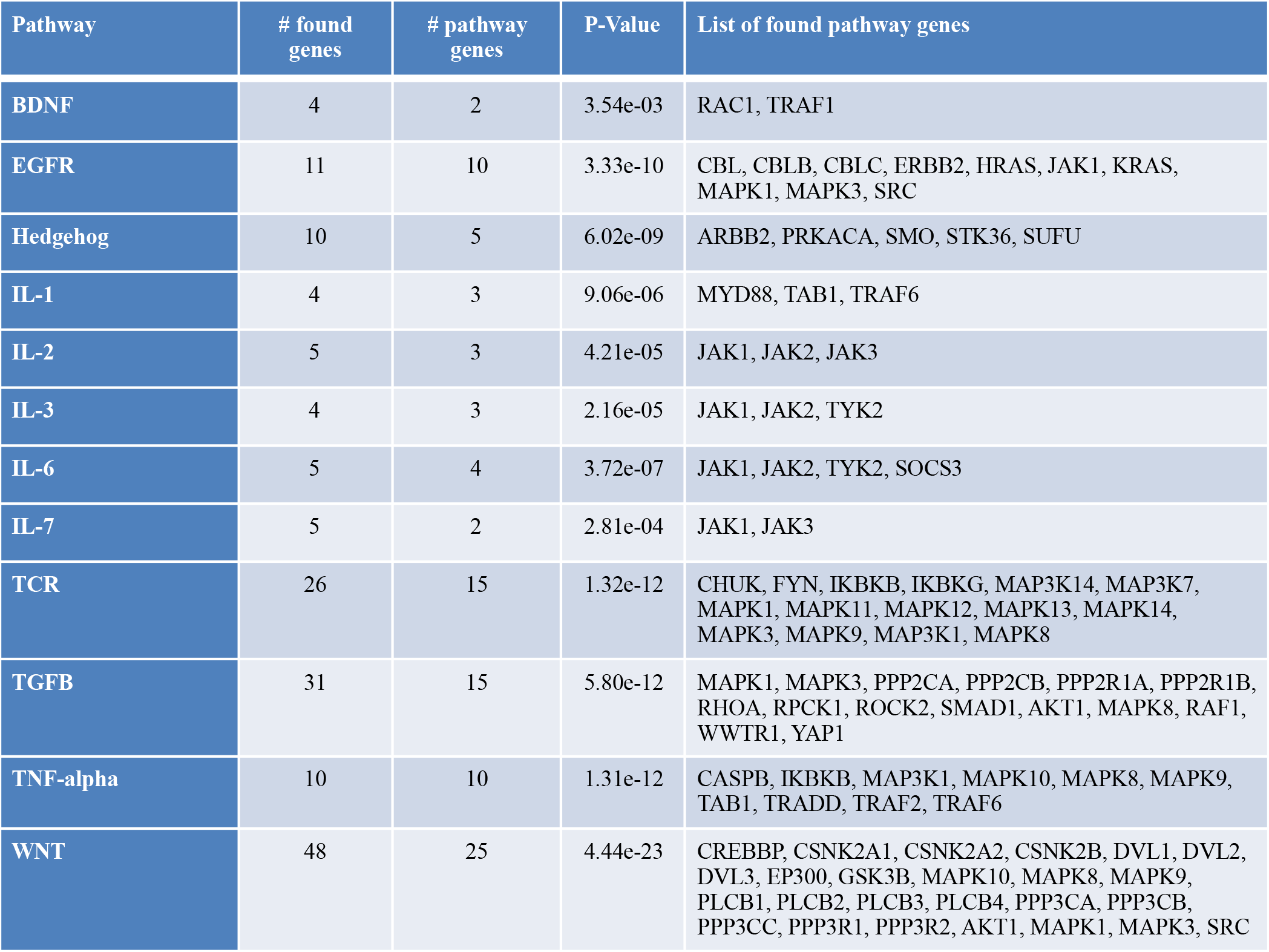
Details of identified strongest path(s) by the application using the KEGG background network

For all pathways, as seen in Table 1, where we used the KEGG network to find the strongest path(s) between receptors and TFs of each pathway, more than 50 percent of genes found by the application were already known as pathway genes in the NetPath database. Also when the STRING network was used, as given in Table 2, approximately 80 percent of genes in the identified strongest path(s), for some pathways 100%, were reported to be pathway genes in the NetPath database. As seen in Table 2, by increasing the threshold parameter and identifying sub-optimal strongest path(s), this amount will decrease.

**Table 2.**
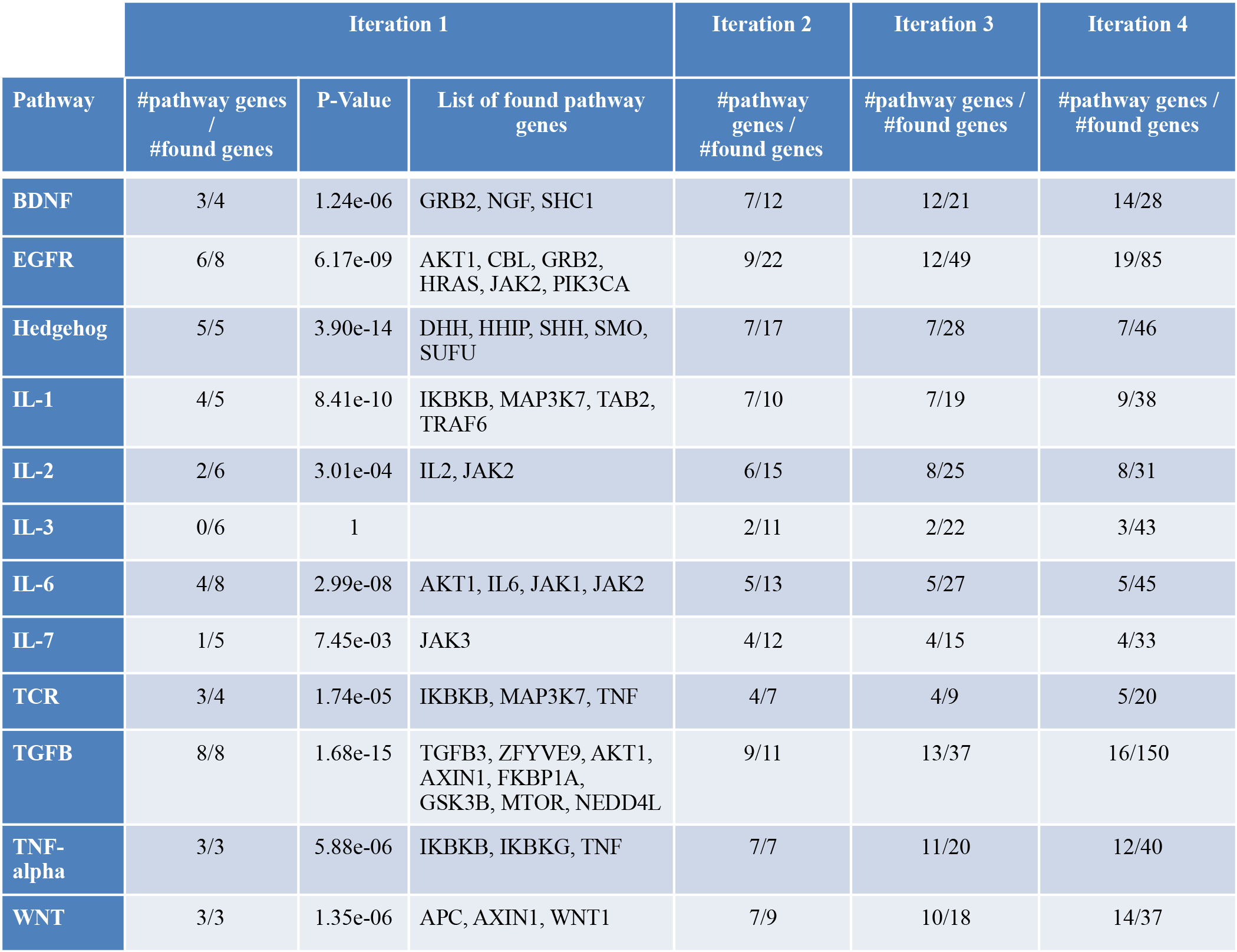
Details of identified strongest path(s) and sub-optimal strongest path(s) by the application using the STRING background network in four separate runs

Our results demonstrate the application can be used for indicating genes in the middle of the pathway by finding strongest path(s) in the signalling network like KEGG, or in the protein interaction network such as STRING. Since the networks are available in the application for both Human and Mouse species, using the StrongestPath app will be very fast and easy compared to similar Cytoscape apps like PathLinker. Although more species can be added to the application in the future without any new installation, currently the application can be used for other species by giving the annotation and the network files to the app manually.

For each pathway, the p-value was calculated by hypergeometric distribution to quantitatively assess the significance of the overlap between the application output and the pathway genes. The p-value measures the probability of detecting the number of genes involved in a given pathway in the middle of the strongest path(s) found by the application. For a given signalling pathway, we define the probability of observing k or more pathway genes in the strongest path(s) containing n genes in the middle as:

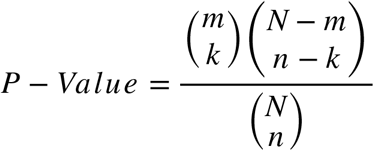

where *N* refers to the total number of genes of the background network (i.e. KEGG or STRING networks in our experiments), and m denotes the number of genes which are already known as pathway genes in the NetPath database. When the p-value is close to zero, it means that most of the genes in the strongest path(s) have already been identified as genes of a given pathway, and it is unlikely that this happens by chance.

According to the obtained results on the above signaling pathways, analyzing the strongest path(s) between source and target genes by considering different PPI or signaling networks as a background network can detect different sets of proteins in the middle of the path, with little in common. Therefore, researchers can easily use the StrongestPath with multiple PPI or signaling networks provided in the application to find which proteins may play a significant role in the middle of the pathway between source and target proteins.

In the following, we only compare the StrongestPath with the PathLinker, which is a Cytoscape application with better accuracy compared to the others for pathway reconstruction, as stated in [4, 5]. The idea of finding the strongest path in the network, used by the PathLinker in 2016, appears for the first time in the first release of our application i.e. StrongestPath in 2015. Both applications also detect sub-optimal paths, but use different approaches to identify and contrary to what is claimed in the PathLinker manuscript [5], the StrongestPath can also identify all sub-optimal paths correctly with a more efficient algorithm. Regarding the same idea used in both applications, the output of both applications is expected to be the same, but there are some benefits, as listed below, which make the StrongestPath more applicable than the PathLinker, and this encourages us to introduce the new release of our application in this manuscript more than before:

1. In addition to allowing users to use their own networks similar to the PathLinker, the StrongestPath makes it possible easily to search for the most probable paths between source and sink proteins in large networks from public databases such as KEGG and STRING. While, when using the PathLinker, you have to load this data manually and in many cases it will be impossible due to the large size of the entire network.
2. Since the number of paths must be given to the PathLinker as an input parameter *k*, users must run the application using different input values to be able to identify the strongest paths that have the same path weight. However, the StrongestPath returns all sub-optimal path(s) with the same path weight in one run. By increasing the threshold parameter in the StrongestPath, the number of nodes in the middle of the network increases, and hence the number of strongest paths increases exponentially. So to reduce the execution time of the application, all paths are not displayed at the output and only all proteins that are at least in the middle of one strongest path are shown.
3. The users can input list of proteins into the StrongestPath via a number of different nomenclatures. So the user doesn’t need to know one specific identifier of their input proteins and in most cases the ID mapping is not necessary to be done before using our application.
4. In the current version of the StrongestPath, identifying regulatory paths (Activatory/ Inhibitory) between transcription factors and target genes can also be done using the TRRUST database.

## Conclusions

In summary, StrongestPath is a Cytoscape application for protein-protein interaction and signaling network analysis. It allows to search for strongest path(s) or sub-optimal strongest path(s) in a PPI or signaling networks for pathway reconstruction, to create and expand network of interactions among a list of proteins and to explore activation or inhibition regulatory paths between TFs and target genes in a regulatory network. Easy access to public large databases and a user-friendly graphical interface make this application more convenient for the users.

## Availability and requirements

### Project name

StrongestPath: a Cytoscape application for protein-protein interaction analysis

### Version

2.0

### Project home page

http://github.com/zmousavian/StrongestPath

### Operating systems

Platform independent

### Programming language

Java

### Software requirements

Cytoscape 3.0 (http://www.cytoscape.org/)

### License

BSD-like license (see website)

## Declarations

### Ethics approval and consent to participate

Not applicable.

### Consent for publication

Not applicable.

### Availability of data and materials

All datasets generated and analysed during the current study are available in the GitHub repository, [http://github.com/zmousavian/StrongestPath]

### Competing interests

The authors declare that they have no competing interests.

### Funding

No funding.

### Authors’ contributions

ZM and ASZ developed the theoretical framework, conceived the idea for the plugin and designed the software architecture. MK, AN and AM implemented StrongestPath, under the supervision of ZM and ASZ. ZM and ASZ supervised the project and coordinated the research team. ZM and ASZ wrote the paper. All authors read and approved the final manuscript.

## Acknowledgements

Authors thank Marcos J. Araúzo-Bravo, Mehrdad Tahvilian and Mehdi Sadeghi for his valuable comments in plugin development and thank Faezeh Shekari for her useful comments about the signaling pathways. We used the computer clusters of the Institute for Research in Fundamental Sciences IPM, Tehran, and Max Planck Institute for molecular biomedicine, Muenster.

## Availability

The StrongestPath application can be downloaded from the Cytoscape App Store (http://apps.cytoscape.org/apps/strongestpath) and the source codes and a user detailed manual are freely available at: (https://github.com/zmousavian/StrongestPath).

## Notes

http://github.com/zmousavian/StrongestPath

http://apps.cytoscape.org/apps/strongestpath

### Competing Interest Statement

The authors have declared no competing interest.

